# The scar constituent Collagen I triggers coordinated collective migration and invasion in a 3D spheroid model of early endometriotic lesions

**DOI:** 10.1101/2020.04.02.005322

**Authors:** Anna Stejskalova, Victoria Fincke, Melissa Nowak, Yvonne Schmidt, Marie-Kristin von Wahlde, Sebastian D. Schäfer, Ludwig Kiesel, Burkhard Greve, Martin Götte

**Author notes:** address correspondence to Prof. Dr. Martin Götte, Department of Gynecology and Obstetrics, Albert-Schweitzer Campus 1, D11, 48149 Münster, Germany, or to Dr. Anna Stejskalova.

## Abstract

Endometriosis is a painful gynaecological condition characterized by ectopic growth of endometrial cells outside of the uterus. Little is known about the mechanisms by which endometrial fragments invade tissues. This is partially due to a lack of suitable experimental models. In this study, we show that a spheroid 3D model, but not single cells mimic the collective endometrial fragment-like invasion through the extracellular matrix. This model reveals that collagen I, the main constituent of surgical scars, significantly increases the rate of lesion formation by healthy endometrial stromal cells (St-T1b) in vitro compared to the basement membrane-like matrix Matrigel. Stromal cell invasion of collagen I requires MMPs, whereas collective migration of endometriotic epithelial 12Z cells involves Rac-signalling. We show that inhibiting ROCK signalling responsible for actomyosin contraction increases the lesion-size. Moreover, endometriotic epithelial 12Z cells, but not eutopic stromal cells St-T1b migrate on Matrigel. The rate of this migration is decreased by the microRNA miR-200b and increased by miR-145. Our 3D model offers a facile approach to dissect how endometrial fragments invade tissues and is an important step toward developing new personalized therapeutics for endometriosis. Moreover, our model is a suitable tool to screen small molecule drugs and microRNA-based therapeutics.

## Introduction

Endometriosis is a common gynaecological disease in which the uterine lining, the endometrium, grows at ectopic locations such as the ovaries and peritoneal cavity^1^. It is currently treated using hormonal therapy and excision surgery. Unfortunately, these treatments are not curative and have high associated side effects and remission rates^1,2^. While targeted therapies are urgently needed, their development has been hindered by the heterogeneity^3,4^ and limited mechanistic understanding of the disease.

Based on the widely accepted Sampson’s theory, endometriosis arises when tissue fragments shed during menstruation implant in the surrounding tissue^5^. In order to implant, endometrial fragments have to first penetrate either through intact barriers consisting of epithelial cells, basement membranes and collagen or directly through surgical scars and then spread^6^. In this regard, endometriosis shares many similarities with metastatic cancer^7^. Nevertheless, while cancer researchers have devoted considerable attention to dissecting the invasive processes involved in cancer metastases^8^, little is known about invasive processes in endometriosis.

A significant hurdle in studying how endometrial cells invade tissues has been a lack of suitable *in vitro* experimental models^9^. Previous *in vitro* models consisted of endometrial explants or single cells combined with chorioallantoic membrane^10^, amniotic membrane^11^, peritoneal mesothelial cell monolayers^12^ and peritoneal explants^13^. However, all of these approaches have some inherent limitations. Explants suffer from high heterogeneity, mixed cell population, low throughput and are confounded by the phase of the menstrual cycle. Single cells are easy to characterize, but their invasive and migratory strategies are markedly different from the coordinated multicellular collective invasion through the extracellular matrix (ECM) that have been observed *in vivo*^14,15^. While endometrial spheroids^16,17^ bear close similarity to endometrial tissue the strategies these multicellular constructs employ to invade the ECM have not yet been investigated.

In this study, we show that endometrial spheroids create structures resembling lesions on collagen I and Matrigel *in vitro* within 5-7 days. We demonstrate that this easy-to-adopt assay can dissect the effect of the cell and ECM type as well as of small molecule- and RNA-drugs. Finally, by focusing on the mechanoregulatory apparatus involved in the development of endometrial lesions rather than on characterizing the eutopic endometrial tissue or mature lesions alone^2^, this study offers a novel approach to studying the pathogenesis of endometriosis.

## Results

### Eutopic stromal and ectopic stromal and epithelial cells self-organize into spheroids in hanging drop culture

A typical lesion consists of both stromal and epithelial endometrial cells^18^ and it is still being investigated whether these cells are clonal in origin and whether the heterogeneity arises due to epithelial-to-mesenchymal transition (EMT)^19^ or the reverse process, mesenchymal-to-eptihelial transition (MET)^20^. To capture the heterogeneity of endometrial cells found in lesions, the cells we employed in this study were an immortalized eutopic stromal cell line St-T1b^21^, primary ectopic endometriotic stromal cells (ESCs) and the ectopic light red peritoneal lesion derived epithelial 12Z cell line that was previously shown to be invasive in a Matrigel invasion assay^18^.

First, we validated that the cells retained their stromal and epithelial morphology in culture. **Figure 1 A** shows that while the St-T1b and ESCs cells have an elongated, fibroblast-like stromal morphology, 12Z cells have a mostly polygonal shape and grow in clusters. Furthermore, on tissue culture (TC) plastic, the stromal cells exhibit more defined actin fibres compared to the 12Z cells. Quantitative analysis (**Figure 1 B**) confirmed that 12Z cells are significantly smaller (p<0.0001) than St-T1b and ESCs, where the average area for St-T1b, 12Z and ESCs cells growth on TC plastic were 2086±904.1μm^2^ (n=29), 787.7± 380.9 μm^2^ (n=32) and 1989±889.5 μm^2^ (n=30).

**Figure 1.**
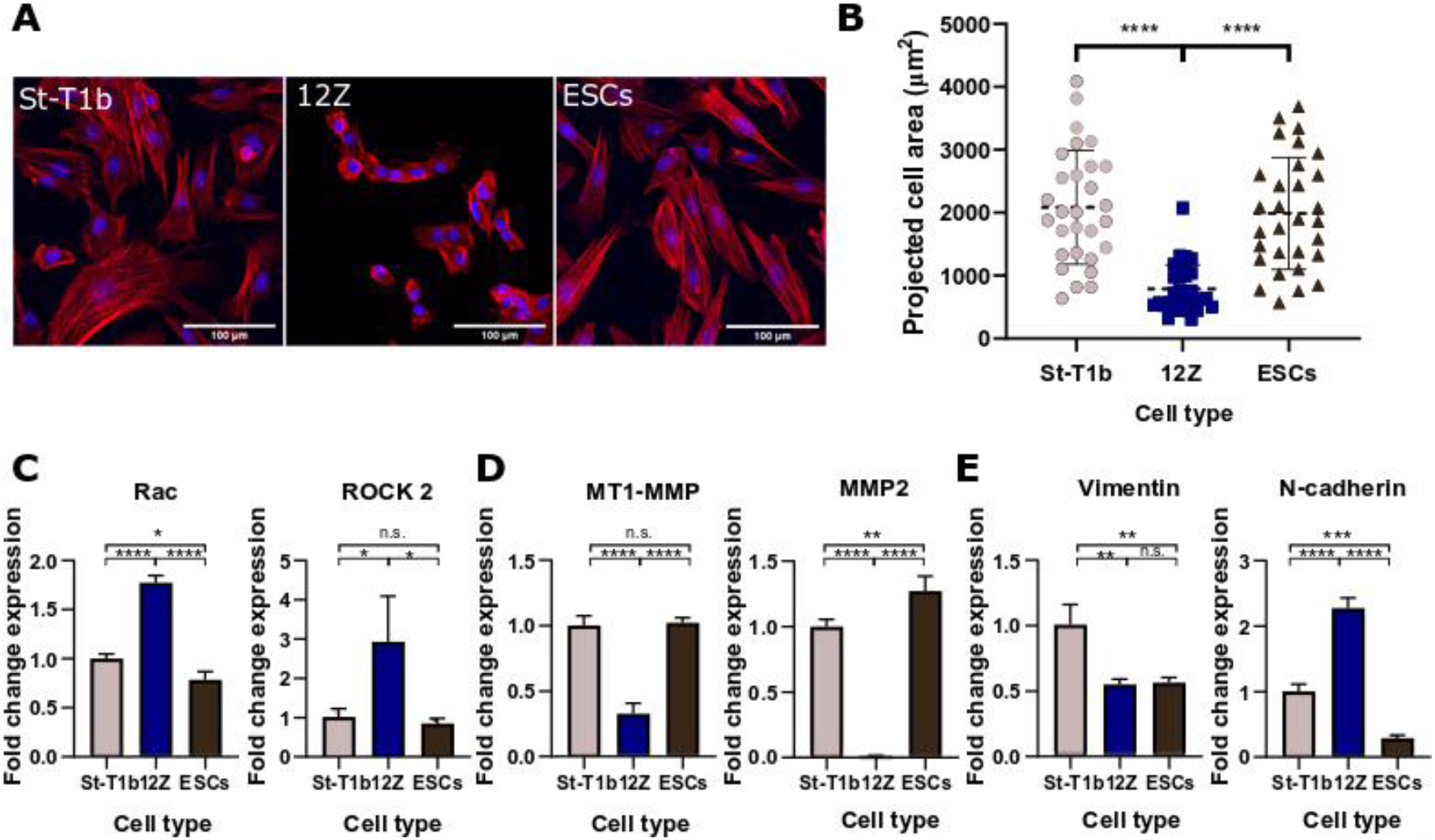
Characterization of endometrial cells (A) St-T1b, 12Z and ESCs on plastic, actin cytoskeleton stained with phalloidin (red) and nuclei with DAPI (blue). St-T1b and ESCs appear spindle-like while 12Z have a cobblestone morphology. Scale bar, 100μm. (B) The projected area in 2D of St-T1b and ESCs is significantly larger than of 12Z cells (n=29, 32 and 30 cells, Kruskal-Wallis with Dunn’s multiple comparisons post-hoc test. Data show mean±s.d.). (C) PCR analysis of Rac1 and ROCK2 shows their overexpression by 12Z cells compared to St-T1b and ESCs (D) MT1-MMP and MMP2 expression is significantly higher in stromal cells compared to 12Z cells (E) The EMT marker vimentin was downregulated in 12Z and ESCs compared to St-T1b and N-cadherin was upregulated in 12Z cells. In all PCR experiments n=3, ANOVA with Tukey’s multiple comparisons post-hoc test, fold change was normalized to St-T1b; data represent mean+s.d. For all figures in this panel *p<0.05; **p<0.01; ***p<0.001, ****p<0.0001 and n.s. p>0.05

Gene expression analysis revealed that each cell type has a distinct signature of invasive markers. First, we examined the expression of Rac1, a small signalling G protein that directs actin-driven cellular protrusion, microtubule prolongation and the formation of lamellipodia^22^ both in single cells and at the leading edge during collective migration^23^. The expression of Rac1 was significantly upregulated in the 12Z cells compared to the healthy St-T1b (p<0.0001, n=3) and diseased ESCs (p<0.0001, n=3) (**Figure 1 C**). The Rho-associated kinase 2 (ROCK2), a kinase that regulates the formation of actomyosin stress fibres and cellular contraction^22^, was significantly upregulated in 12Z cells compared to St-T1b and ESCs (p=0.0338 and p=0.0237, respectively, n=3).

Stromal cells exhibited higher proteolytic gene expression compared to epithelial cells (**Figure 1 D**). qPCR analysis revealed that the stromal cells exhibit higher expression of MT1-MMP (p<0.0001, n=3) and MMP-2 (p<0.0001, n=3) compared to epithelial cells. The expression of MMP-1 and MMP-7 was not detectable in any of the studied cell types.

As the EMT/MET processes have been implicated in the progression of the disease, we further investigated the expression of mesenchymal markers vimentin and N-cadherin (**Figure 1 E**). Vimentin was downregulated in both the 12Z and ESCs cells compared to the control St-T1b cell line (p=0.0024 and p=0.0029, respectively, n=3). The expression of N-cadherin, a cadherin known to promote invasion in many cell types^24^, was upregulated in the 12Z (p<0.0001, n=3) cells and downregulated in ESCs (p=0.0006, n=3) compared to St-T1b cells.

#### Eutopic stromal, ectopic stromal and epithelial cells self-organize into spheroids in hanging drop culture

Recent studies suggested that spheroid culture offers several advantages over 2D culture and confirmed that 12Z cells^16^ and endometriotic stromal cells^17^ can assemble into spheroids using the U-bottom 96 well plates^17^. However, it has not been investigated whether also the hanging drop method can be used to fabricate endometrial spheroids and whether there are any differences between spheroids formed from epithelial and stromal endometrial cells. We, therefore, evaluated the hanging-drop method, each drop containing 20 000 of either stromal or epithelial cells in 20μl of standard media and selected day 4 as the harvesting day.

Bright-field images (**Figure 2A**) show that all the studied cell types self-organized into spheroids. Interestingly, the morphology of the spheroids varied across cell types. St-T1b and ESCs cells assembled into compact, round-spheres, while the 12Z spheroids were larger and sometimes exhibited slightly branching morphology. To quantify this, circularity and area of the spheres were analyzed. The circularity values were 0.956± 0.027 (n=11) for the St-T1b cells, 0.944± 0.067 (n=11) for ESCs, and 0.890± 0.095 (n=11) for 12Z cells and did not vary significantly (p>0.05) across the studied cell types (**Figure 2B**). Interestingly, while the size of individual 12Z cells in 2D is significantly smaller compared to the ESCs and St-T1b cells, 12Z spheroids were significantly larger compared to St-T1b and ESCs (n=14, p<0.0001 and p<0.001) (**Figure 2C**). In order to exclude that this is due to a cell-counting error, the spheroid size was measured on spheroids prepared three independent times.

**Figure 2.**
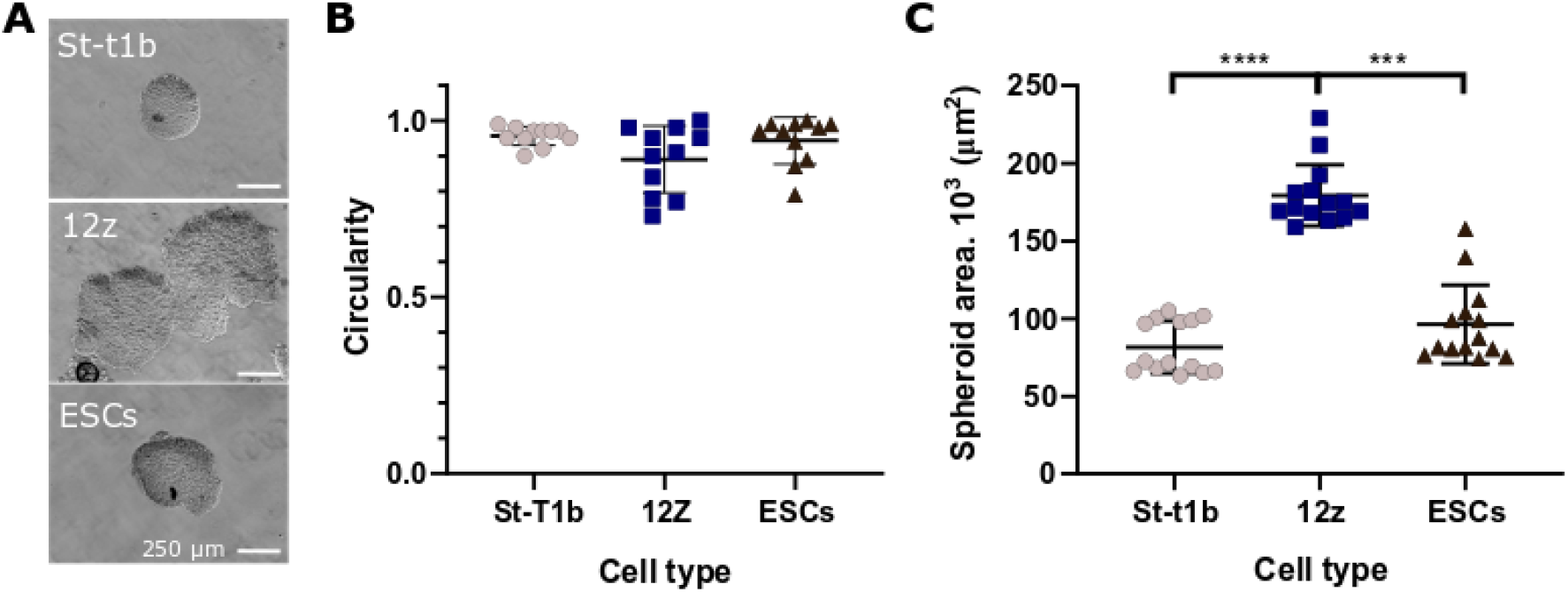
Spheroid formation by endometrial cells. (A) Bright-field images of fixed spheroids that formed after 4-days using the hanging drop method. Scale bars.250μm. (B) All spheroids prepared across three preparations were circular (n=11, Kruskal-Wallis, n.s.) (C) The 12Z spheroids were significantly larger compared to St-T1b and ESCs spheroids (n=14, Kruskal-Wallis with Dunn’s multiple comparisons post-hoc test), the area was measured manually on bright-field images, 10x magnification; For all figures in the panel *p<0.05; **p<0.01; ***p<0.001, ****p<0.0001 and n.s. p>0.05;

### Spheroid 3D culture reveals that endometrial cells adopt a range of invasive and migratory strategies that are modulated by the ECM

Having confirmed that both stromal and epithelial endometriotic cells were able to form spheroids, we evaluated their invasive behaviour on two different ECM-derived hydrogels–Matrigel and collagen I.

On the basement membrane (BM) mimic Matrigel, the stromal St-T1b spheroids remained rounded with ESCs exhibiting few protrusions and only the 12Z spheroids consistenly developed multiple multicellular protrusions across several preparations. Confocal imaging (**Figure 3A**) revealed that the 12Z protrusive edges consisted of tightly packed cells (DNA in blue) with scant cytoplasm (actin staining in red). The invading structures emanated not only from the spheroid edges, but individual invasive branches could also be discerned throughout the spheroids on optical sections obtained by confocal microscopy (**Figure 3F**). This suggests Matrigel signalling triggers sophisticated cell-cell communication and coordination within the spheroids. The behaviour of single cells on Matrigel was similarly cell-type dependent (**Figure 3B, top**). The St-T1b and ESC single cells on Matrigel (**Figure 3B**) self-organized into rounded cellular aggregates and 12Z single cells organized into protrusive aggregates.

**Figure 3.**
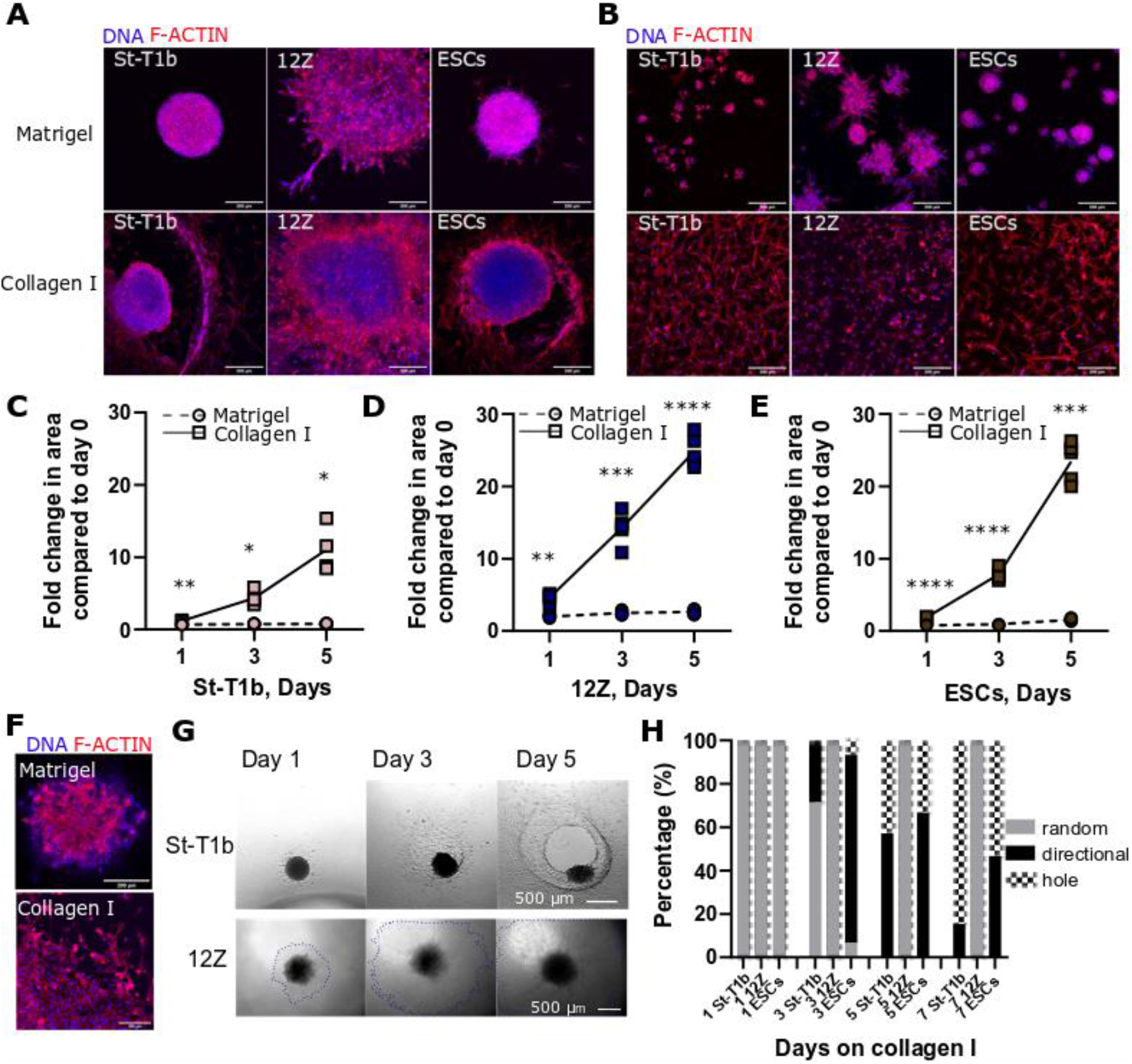
Lesion-like structures on collagen I and Matrigel. (A) Confocal images of spheroids after 7 days on Matrigel and collagen I. 12Z exhibited the highest number of protrusions on Matrigel. St-T1b and ESCs created circular defects on Collagen I surrounded by cells, whereas epithelial 12Z cells migrated as a sheet and confocal imaging revealed no invasion (maximal intensity projection, scale bar, 200μm, actin cytoskeleton red, nuclei blue). (B) Confocal images of a suspension of endometrial cells after 3 days on Matrigel (top row). Stromal St-T1b and ESCs cellular aggregates consisted of only a few cells and were highly circular and 12Z aggregates were larger and showed protrusions. All cell types invaded collagen I (bottom row) as single cells. (maximal intensity projection, scale bar 200μm, actin cytoskeleton red, nuclei blue) (C) Quantification of St-T1b spheroid growth on collagen (squares, full line) compared to Matrigel (circles, dashed line) (n=4-5, two-way Repeated Measures (RM) ANOVA, Šidák’s multiple comparisons test), (D) Quantification of 12Z spheroid growth on collagen I (squares) compared to Matrigel (circles) (n=5, two-way RM ANOVA, Šidák’s multiple comparisons test) (E) Quantification of ESCs spheroid growth on collagen (squares) compared to Matrigel (circles) (n=5, two-way RM ANOVA, Šidák’s multiple comparisons test),(F) Top: Confocal optical section through the middle region of a 12Z spheroid on Matrigel on day 7 shows branching of the spheroid. Bottom: Edge of 12Z cells spreading on collagen I. Cells engage in collective migration with a few individual cells breaking away. Scale bar 200μm, (G) Stromal St-T1b spheroids first exhibited random spreading, followed by a directional invasion and defect formation. Spreading of 12Z cells on collagen proceeded evenly in all directions. Scale bar, 500μm (H) Quantification of lesion shapes on collagen I over 7 days. Data were compiled from three independent experiments (n=13-15 per time point). For all figures in this panel *p<0.05; **p<0.01; ***p<0.001, ****p<0.0001, and n.s. p>0.05;

The response of all studied cell types both as spheroids and single cells was markedly different on the scar constituent collagen I^25^ compared to Matrigel **(Figures 3A-E)**. St-T1b and ESC spheroids on collagen I developed into invasive lesion-like structures **(Figures 3A)**. More specifically, the St-T1b and ESC spheroids gradually invaded collagen I, leaving behind a defect with a ring of tightly adhering cells at its margins. These rings appeared to stabilize the defect and to limit further random cellular spreading outside of the defect in many but not all spheorids. In order to understand better the coordinated spreading of spheroids on collagen, we quantified the invasive and migratory patterns on this ECM component using bright-field imaging. The stromal cells (St-T1b and ESCs) first exhibited random spreading **(Figure 3G-H)** followed by directional spreading by day 5. Finally, a defect formed in the area with the densest stromal cell population, leaving behind a hole. In our system (3mg/ml, 40μL/well) this typically occurred around day 5 or 7 with 84.6% and 53.3% of St-T1b and ESCs having a defect on day 7(n=13-15 per time point) (**Figure 3H**).

Unlike the stromal cells, the 12Z spheroids on collagen I did not form any defects and engaged in a rapid collective surface migration as confirmed by microscopy (**Figure 3A, D, G, H**). We could further observe that some cells close to the leading edges separated and migrated as single cells (**Figure 3A** and **3F**).

In contrast to spheroids on collagen I, single cells on collagen I did not engage in coordinated invasion and invaded collagen I as single cells without any apparent cell-cell interactions (**Figure 3B**). Overall, regardless of cell type, the combined migration or invasion of spheroids, quantified as a projected surface area, was significantly more prominent on collagen I compared to Matrigel when quantified on days 1, 3 and 5 (**Figures 3C-E**). In summary, these results highlight the importance of ECM type, cell type and cell-cell contact in endometriosis pathology.

### Spheroid 3D culture as an effective tool to screen small molecule drug and microRNA-based therapeutics

Having established that spheroids on various ECM components functionally mimic many of the predicted invasive phenotypes observed histologically i*n vivo*^14,15^, we used this assay to study the mechanisms involved in the spreading of early lesions and the potential therapeutic effect of mechanoregulatory small molecules and micro RNAs.

#### The broad-spectrum MMP inhibitor NNGH limits the invasive behaviour of stromal but not epithelial endometriotic spheroids on collagen I

Previous studies implicated MMP signalling in the formation of early endometriotic lesions^10^. Our study shows that the broad-spectrum MMP inhibitor 15μM N-isobutyl-N-(4-methoxyphenylsulfonyl) glycyl hydroxamic acid (NNGH) significantly reduced the spreading and invasion through collagen from 15.27 fold to 2.05 fold (n=4, p=0.0087) and 10.81 fold to 3.61 fold (n=5, p<0.001) in St-T1b and ESCs, respectively, but did not significantly affect the spreading in 12Z cells (n=3-5, n.s.) (**Figure 4A**). Furthermore, it can be seen from **Figure 4B**, that while NNGH treatment prevents the formation of the defect on Collagen I even after 7 days in culture, the migration of St-T1b and ESCs is not completely eliminated.

**Figure 4.**
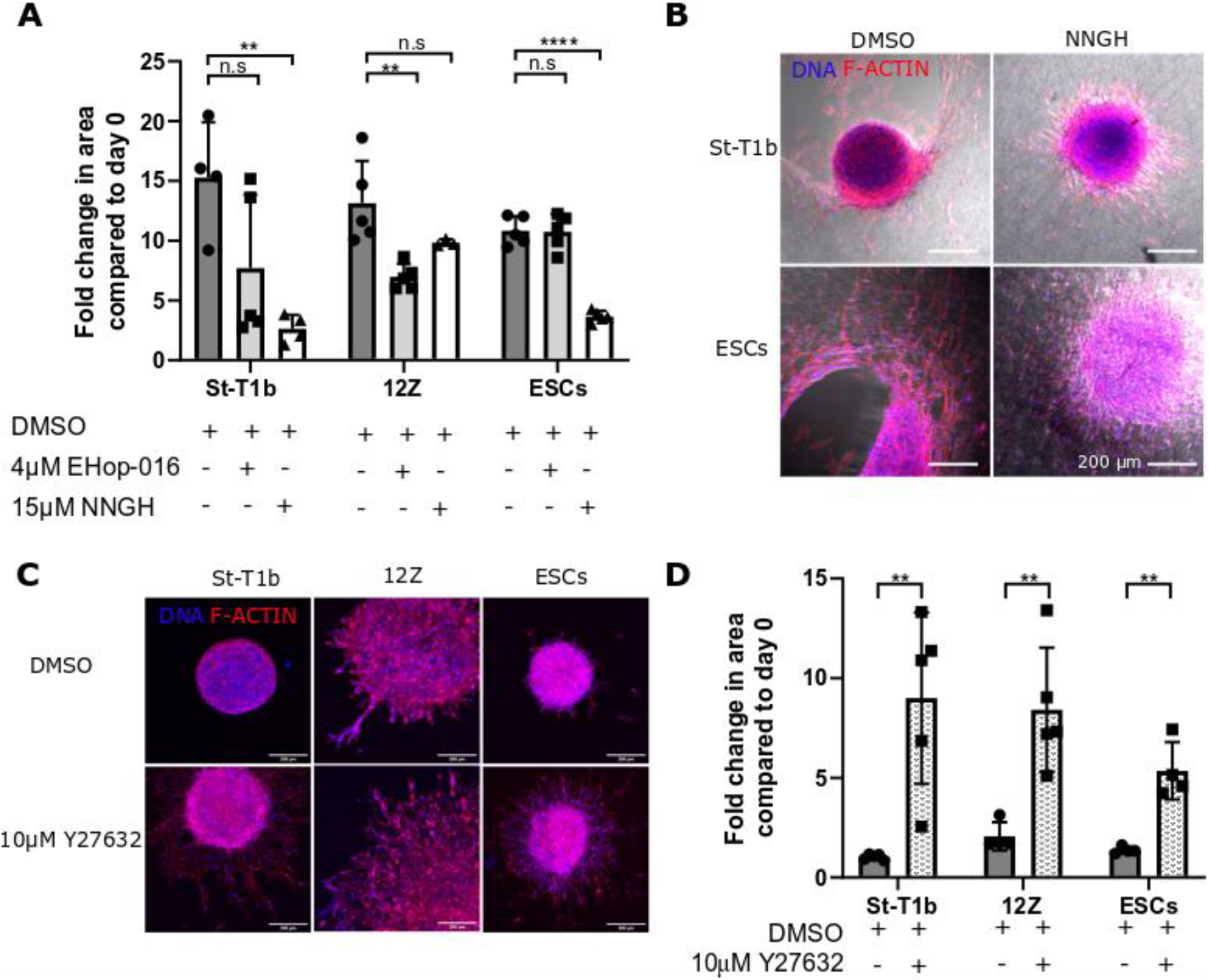
The effects of small molecule inhibitors on lesion formation. (A) The broad spectrum NNGH inhibitor significantly reduced the in vitro lesion size in St-T1b and ESCs but not in 12Z cells that migrated on collagen I surface. The Rac inhibitor EHop-016 significantly reduced the lesion size in 12Z cells, but not in St-T1b and ESCs. EHop-016 affected viability in St-T1b cells. The spheroid size was measured manually on days 0 and 5 (n=3-5, ANOVA with Tukey’s multiple comparisons post-hoc test was performed separately for each cell type to compare treatments) (B) The broad-spectrum MMP inhibitor NNGH effectively prevented stromal cells from degrading collagen I (bright field channel) but did not completely prevent the cells from migrating. Confocal images were obtained on fixed samples after 7 days in culture. Scale bar, 200μm (C) ROCK inhibitor Y27632 significantly promoted spreading on Matrigel even in St-T1b. Confocal images were obtained on fixed spheroids on day 7, f-actin in red and DNA is stained in blue. Scale bar, 200μm. (D) Y27632 significantly increased the spreading of endometrial cells on Matrigel after 5 days. Data were compared to the spheroid size on day 0 using bright-field images (n=5, multiple t-tests). *p<0.05; **p<0.01; ***p<0.001 and n.s. p>0.05;Data shown as mean± s.d.,

#### The Rac inhibitor EHop-016 reduces the rate of 12Z collective migration on Collagen I

Next, we evaluated the effect of the Rac inhibitor EHop-016 at 4 μM on the spreading of endometrial cells on collagen I (**Figure 4A**). The Rac inhibitor significantly reduced the invasive area in 12Z cells from 13.14 to 6.97 (n=5, p=0.005) after 5 days on collagen I. Moreover, the Rac inhibitor also affected the invasive behaviour and viability of some St-T1b spheroids on collagen I.

#### ROCK inhibition significantly enhances spreading and invasion of all studied endometrial cell types

The ROCK inhibitor Y27632 significantly (p<0.01) increased the spreading of all studied cell types on Matrigel (**Figure 4C**). The area occupied by St-T1b, 12Z and ESCs was 9, 8.41 and 5.35 fold larger following Y27632 compared to DMSO treatment alone after 5 days in culture (**Figure 4D**).

#### The spheroid model reveals context-dependent roles of the mechanoregulatory microRNAs miR-200b and miR-145 on the invasive behaviour of endometriotic epithelial cells on Matrigel

We next investigated whether our *in vitro* model can be used as a high-throughput tool to evaluate the functional effect of various microRNAs on the early stages of endometriosis. In particular, we selected two microRNAs, miR-200b^26^ and miR-145^27^, that have been previously shown to be dysregulated during endometriosis^28^ and to modulate the invasion and migration of 12Z cells in 2D and Transwell assays. miR-200b acts as a transcriptional repressor of ZEB1/2 and thus downregulates EMT transition by suppressing the expression of Vimentin and promoting the expression of E-cadherin^29^. The miR-145 is upregulated in endometrial lesions and has been described to modulate cytoskeletal dynamics in a number of cell types, including endometrial, and has a number of validated targets, including beta and gamma actin, cofilin, fascin, myosin light chain 9 and Rho kinase Rock1^27,30^. The transfection was performed in monolayer culture prior to the fabrication of spheroids and the effect of microRNAs on spheroid growth was assessed after 3 days on Matrigel to minimize the effect of miR dilution and degradation^31^ (**Figure 5A**). It can be seen from **Figure 5B** that microRNA transfection did not significantly alter the ability of cells to form spheroids and the area of individual spheroids was not significantly different (p>0.05) across the treatment groups. The microRNAs, however, affected the invasive behaviour of 12Z cells on Matrigel as seen in the bright-field images in **Figure 5C**.The epithelial phenotype-promoting microRNA miR-200b significantly decreased the number of sprouts per spheroid from ~19 to ~1 (p=0.0002) while miR-145 significantly increased the number of sprouts per spheroid to ~35 (p=0.0005) (**Figure 5D**) and increased the overall sprouting area from 71.13·10^3^± 14.54·10^3^ μm^2^ per scrambled control miR spheroid to 158.06·10^3^±24.67·10^3^ μm^2^ per miR-145 treated spheroids (p<0.0001) (**Figure 5E**).

**Figure 5.**
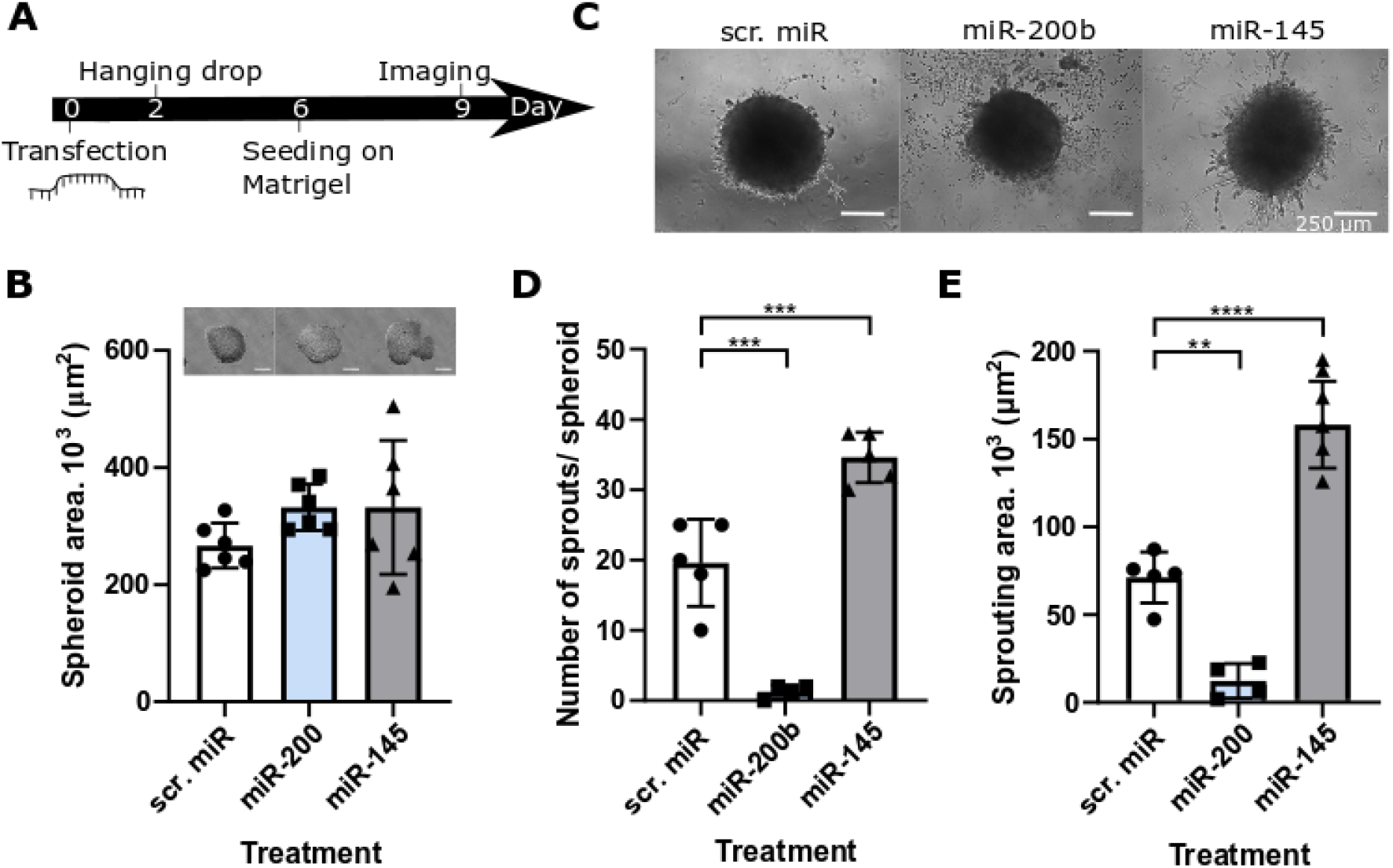
The effect of microRNA on 12Z spreading on Matrigel. (A) Workflow (B) none of the microRNAs affected the ability of 12Z cells to self-organize into spheroids (scale bar=250μm, n=6, ANOVA), (C) Representative images of microRNA treated 12Z spheroids after 3 days on Matrigel. Scale bar, 250 μm. (D) miR-200 significantly decreased while miR-145 significantly increased the number of sprouts per spheroid after 3 days on Matrigel. (n=4-5, ANOVA, Tukey’s multiple comparisons) (E) The overall area occupied by sprouts was significantly larger and smaller when treated with miR-145 and miR-200b, respectively, compared to scr.miR after 3 days on Matrigel (n=4-5 randomly chosen spheroids, ANOVA, Tukey’s multiple comparisons), *p<0.05; **p<0.01; ***p<0.001 and n.s. p>0.05; experiments were conducted two independent times, data expressed as mean±s.d.

## Discussion

The overall goal of this study was to develop a modular, multicellular 3D *in vitro* system that makes it possible to study the interplay of different factors affecting the pathogenesis of the early endometriotic lesion formation *in vitro*.

We demonstrate that the hanging drop method makes it possible to generate spheroids of reproducible size and thus provides a good alternative to the low-adhesion plate method^16^, ^32^. Our data show that the spheroid size is consistently cell-type specific, with stromal cells generating more compact spheroids than the smaller epithelial 12Z cells.

Our spheroid model revealed that endometriotic cells employ a host of invasive and migratory strategies that are cell type and ECM dependent. In particular, the invasive modes cells adopted using the 3-D spheroid model but not single cells on ECM could, for the first time, reproduce collective-migration phenotypes previously identified only through histological and clinical observations, further confirming the relevance of this model.

Several clinical studies suggested endometriosis can only implant in damaged tissues^33,34,35^. To investigate this *in vitro*, we mimicked the intact tissue using the basement membrane mimic Matrigel and scar-tissue mimic collagen I. All cell types displayed markedly different phenotype and a significantly higher level of spreading on collagen I compared with Matrigel, suggesting a profound role of the ECM in modulating the growth of early lesions. On Matrigel, only ectopic 12Z spheroids consistently exhibited cellular spreading. While 12Z cells could spread both on Matrigel and collagen I, they migrated significantly faster on collagen I. Strikingly, collagen I triggered invasive behaviour not only in ectopic ESCs but also in eutopic St-T1b, stromal cells. Invasion of collagen I was highly orchestrated. The fact that the migration and invasion in St-T1b and ESCs was in all instances directional suggests sophisticated cell-cell communication within these spheroids. The mature *in vitro* lesions that typically developed by day 7 consisted of a collagen defect surrounded by an interconnected cellular ring, resembling a cellular structure previously described by Martin and colleagues in the context of wound healing^36^. Taken together, our data suggest that while patient-derived cells are more likely to penetrate intact tissue compared to those derived from healthy-donor-derived cells, exposed collagen I alone triggers invasive phenotype even in healthy stromal cells. These observations agree with previous studies conducted on cancer cells suggesting that collagen content alone can increase the invasive cellular phenotype^37,38^. In conclusion, our results provide clear support for the clinical data suggesting that endometriosis commonly implants in scars^34^, including surgical scars such as that left behind by the C-section^33^.

We next show that the collagen I defects caused by stromal cells arise due to matrix degradation via MMPs rather than due to cellular contraction and traction forces. Both eutopic and ectopic stromal cells had significantly upregulated MMP expression and the MMP inhibitor NNGH significantly reduced the size of *in vitro* stromal lesions on collagen I. These results are in good agreement with Nap and colleagues that demonstrated that inhibiting MMP activity prevents the development of endometriotic lesions in a chicken chorioallantoic membrane model^10^. Our results refine this model and show that while, in agreement with the previous studies^39^, the MMP inhibitor significantly slowed down the invasion of spreading of stromal cells on collagen it had little effect on the collective migration of ectopic epithelial 12Z cells, suggesting personalized therapeutic strategies and lesion stratification might be necessary. Such a cautious approach is further justified by a recent report of failed MMP inhibitors clinical trials in cancer, despite showing promising preclinical results^40^. Rac inhibitor EHop-016 slowed down, but not inhibited, collective migration on collagen I in 12Z cells and affected cellular viability in stromal cells. A similar cell-type dependent effect of EHop-016 inhibitor on cell viability was previously reported by other studies^41^. Contrary to our expectations, ROCK inhibitor Y27632 that prevents hydrogel contraction in several cell types^42, 43^, significantly increased cellular spreading *in vitro* in all studied cell types. Our results, therefore, provide strong evidence against using ROCK inhibitors for the treatment of endometriosis-associated fibrosis as previously suggested by Yuge and colleagues^44^ and cancer due to the off-target effects on endometrium^45^. Indeed similar context-dependent effects of ROCK inhibitors have been described in microvascular endothelial cells^46^, retinal pigment epithelial cells^47^ and osteoblastic cells^48^.

While the exact cause of endometriosis remains unclear, several recent studies linked the pathology of endometriosis to dysregulated microRNA signaling^28,49^. Given that a typical micro-RNA has tens of targets, sequencing studies need to be accompanied by reliable functional assays to be biologically meaningful^50^. In this study, we demonstrate that the spheroid assay can be used to reproducibly evaluate the effect of individual microRNAs on the complex, multicellular spreading of endometriosis-mimicking constructs over several days. We show that reverting the 12Z phenotype to an epithelial-state using the microRNA miR-200b as we previously demonstrated in our laboratory^26^ significantly reduces spheroid spreading on Matrigel. These results align well with the widely accepted concept that EMT is required for the epithelial cells to migrate on and invade the basement membrane^51^ and our in vitro model thus supports the notion that the EMT/MET transition could be targeted for therapeutic purposes in endometrial patients^52^. Our spheroid model further revealed that, contrary to previous *in vitro* 2D assays^27^, the microRNA miR-145 up-regulated in ectopic lesions *in vivo* increases 12Z spreading *in vitro*. Overall, we demonstrate that the spheroid assay can be used to screen for both small molecule and RNA-based therapeutics (**Figure 6**).

**Figure 6.**
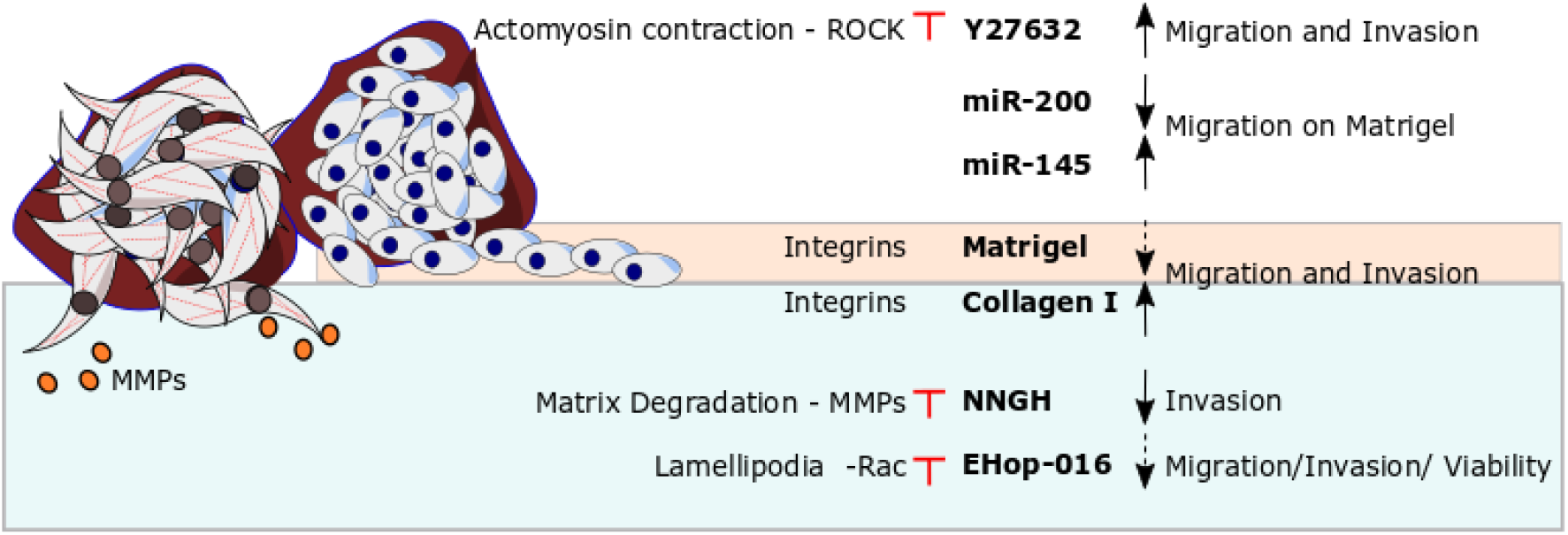
The invasiveness of endometrial spheroids depends both on the cell type (stromal St-T1b and ESCs in brown, epithelial 12Z in blue) and ECM. Invasion and spreading is strongly enhanced by exposed collagen I. Migration on Matrigel is modulated by microRNAs and ROCK inhibitor increases invasion and migration on this substrate. Invasion on collagen is MMP-dependent while collective migration on collagen I involves Rac signalling. Red marks signify experimental intervention. Blunt arrow signifies that an inhibitor was added. Black arrows show the effect of the intervention. Dashed arrow suggest weak effect.

Researchers have recognized for decades that metastatic cells and cells during wound healing possess a remarkable ability to temporarily alter their phenotypes to achieve migration or invasion^53,54^. However, endometriosis has been mostly modelled and studied as a mature and static disease^55^. By focusing on modelling of the invasive processes at ectopic locations in endometriosis, rather than recreating full endometrial tissue *in vitro^56^*, our study presents a new paradigm in *in vitro* endometriosis modelling. Taking into consideration that endometrial cells momentarily change their phenotypes, paves the way for the discovery for new druggable targets.

Future studies will refine the model by including a mesothelial layer and expanding the number of donors and disease subtypes. Furthermore, the processes responsible for the coordinated directional spreading of endometrial stromal spheroids will be explored. Finally, the role of other cells types, such as the immune cells^57,2^ will be investigated.

In conclusion, this study introduces the spheroid assay as an effective, easy to implement method to study the early stages of tissue invasion by endometrial cells and demonstrates good agreement with *in vivo* findings. We anticipate that this assay will be used to gain further insights into invasive processes involved in endometriosis, for high-throughput screening of both small molecule and RNA-based drug candidates and their off-target effects as well as in developing personalized treatment strategies for individual subtypes of the disease.

## Methods

### Cell Culture

The 12Z ectopic epithelial cell line^17^ was maintained in DMEM media (Sigma-Aldrich, cat. No. D0819, Deisenhofen, Germany,) supplemented with 10% FBS (Biochrom GmbH, cat. no. S0615, Berlin, Germany) and 1% Pen/Strep (Sigma-Aldrich, cat. No. P4333). The St-T1b cell line^21^ and primary endometriotic ectopic lesion-derived stromal cells (ESCs) were maintained in 70% DMEM/ 18% MCDB 105 media (Sigma-Aldrich, cat. No. 117-500) supplemented with 10% FBS, 1% Pen/Strep, 1% Glutamine and 5μg/ml insulin (Sigma-Aldrich, cat. No. 10516). Cells were routinely split twice a week. ESCs were prepared from ectopic lesions and characterized as previously described^58^. Primary endometriotic stroma cells were prepared from a biopsiy of a woman with endometriosis who underwent surgical treatment at the Department of Gynecology and Obstetrics of Münster University Hospital in 2013. The modified American Society for Reproductive Medicine classification was used to assess endometriosis^59^. For all ESC experiments, stroma cells derived from a lesion located at the pelvic wall (rASRM score II) of a 19 year old patient were employed. The study was carried out in accordance with the Declaration of Helsinki and approved by the local ethics commission (Ethikkommission der Ärztekammer Westfalen-Lippe und der Medizinischen Fakultät der WWU; approval no. 1 IX Greb 1 from 19 September 2001, updated 2012). The participant gave written informed consent.

### Spheroid formation

Spheroids were generated using the hanging drop method^60^, where 20μL drops each containing 20 000 cells were deposited on the top lid of a plastic Petri dish and the bottom chamber was filled with sterile water or PBS (Sigma-Aldrich, cat. No. D1408). The spheroids were harvested after 4 days at 37°C and 7.5% or 5% CO_2_.

### Preparation of collagen I and Matrigel

A 3mg/ml collagen I hydrogel was formed by neutralizing and diluting the stock solution of Collagen Type I Rat Tail matrix (Corning, Bedford, MA, USA, cat. No. 354236, 4mg/ml or 3.4. mg/ml batch) with 1N NaOH (Applichem, cat. No. A1432, Darmstadt, Germany), 10x PBS (Sigma-Aldrich, cat. No. D1408) and chilled deionized water. The amount of 1N NaOH was calculated as 1N NaOH volume = (volume of the stock collagen) · 0.023 ml. The amount of 10x PBS was calculated as volume 10x PBS = (final volume)/10. Phenol red-free Basement Membrane Matrix Growth Factor Reduced Matrigel (Corning, cat. No. 356231) was used as received and was thawed on ice. The gels were deposited into pre-chilled 96-wells at 35-40 μL per well. Each 96-well plate was subsequently sealed with parafilm and the gels were left to solidify for 30-60 minutes at 37°C. For higher magnification confocal imaging, collagen and Matrigel were deposited on glass coverslips.

### Spheroid response to collagen I/Matrigel

Following gel formation, the wells in a 96-well plate were filled with 50 μL of phenol-red free DMEM (Gibco, cat. No. 21063-029, Darmstadt, Germany) supplemented with 5% charcoal-treated FBS (Biochrom GmbH, cat. no. S0615) and 5μg/ml insulin solution (Sigma-Aldrich, cat. No. 10516). Subsequently, one to three spheres per well were manually added to individual wells. The media were changed every three to five days and the samples were kept in an incubator at 37°C and 7.5% CO_2_. The spheroids were imaged on day 1, 3, 5 and 7.

### Inhibitors

The effects of three inhibitors on spheroid growth were evaluated. The MMP inhibitor NNGH (Merck, cat. No. SML0584, Darmstadt, Germany) was stored at 15mM in DMSO and dissolved to the final concentration of 15μM in media, the Rac inhibitor EHop-016 (Sigma-Aldrich, cat. No. SML0526, stock 4mM, DMSO) was used at the concentration 4 μM and the ROCK inhibitor Y27632 (Sigma-Aldrich, cat. No. Y0503,10mM stock) at 10 μM. In all experiments, spheroids were added directly to inhibitor-containing media. Inhibitor-containing 5% charcoal-treated FBS/insulin media were exchanged every 3 days.

### microRNA transfection

The transfection with negative control microRNA (Scr. miR), miR-200b and miR-145 (Table 1) was performed in a 6-well plate on 60-70% confluent cells. Prior to transfection, the growth media were exchanged for Opti-MEM I Reduced Serum Media (Gibco^®^, cat. no. 31985-070, Thermo-scientific, Germany). The transfection with 20nM microRNA of interest (Table 1) was conducted using the Dharmafect reagent (Dharmacon™, cat. no. T-2001-03, Lafayette, CO, USA). The cells were incubated with the transfection mixture for 24 hours when the media were exchanged for full growth media. MiR spheroids were fabricated 48 hours after the addition of transfection media.

**Table 1.** MicroRNAs used in the study

| MiR | Specifications | Cat. Number | Manufacturer |
| --- | --- | --- | --- |
| Scr. miR | Pre-miR Negative Control 2 | AM17111 | Ambion,<br>Darmstadt,<br>Germany |
| miR-200b | hsa-miR-200b-3p: MC 10492, mirVana, miRNA mimic | 4464066 | Ambion |
| miR-145 | hsa-miR-145, Pre-miR miRNA Precursor | AM17100 | Ambion |

### Staining and immunostaining

The F-actin cytoskeleton was visualized using Phalloidin CruzFluor™ 594 Conjugate (Santa Cruz Biotechnology, cat. No. sc-363795, Santa Cruz, CA, USA) at 1:1000 dilution. The nuclei were visualized using DAPI (Sigma-Aldrich, cat. No. D9564) diluted at 1:50,000. The cells were fixed using 3.7% formaldehyde (Merck, cat. No. 1.04003.1000, Darmstadt, Germany) at 37°C for 15 minutes. Following washing with PBS (Sigma-Aldrich, cat. No. D1408), the cells were permeabilized with 0.1% Triton-X (Riedel-de-Haen, cat. No. AG 56029, Seelze, Germany) for 5 minutes. Hydrogels in a 96-well plate were stained by adding 25μL of the 1:1000 phalloidin dye and incubated for 1 hour at 37°C.

### Imaging

Cells were analysed for morphological and cytoskeletal markers. The bright-field images were obtained using either an Axiovert100 (Carl Zeiss, Jena, Germany) or an inverted microscope (Leica, Wetzlar, Germany) using 5x, 10x and 20x objectives. Confocal imaging was performed on fixed stained samples in a 96 well plate. Samples were imaged with the Zeiss LSM 880 inverted confocal microscope (10x, 0.45 NA) (Carl Zeiss, Jena, Germany) equipped with ZEN 2 software and using 11.04 μm z-stack intervals and sequential scanning (514 nm argon laser, 405 nm diode laser, Bright field). The number of sections was adjusted based on the sample thickness.

### Image analysis

All images were analysed in FIJI^61^. Confocal images are depicted as maximal intensity projections. The spheroid size was measured manually on Bright-field images of spheroids on Petri Dishes, glass slides or in a 96-well plate. Fold increase in area was calculated as the spheroid area on a given day divided by spheroid size on day 0 or day 1 after excluding all the protrusions. The circularity of spheroids was analysed in FIJI that calculates circularity as circularity = 4· π · (area/perimeter^2). The number of sprouts per image was counted manually and the sprouting area was calculated as the total area occupied by an expanding spheroid with sprouts minus the area occupied by a spheroid alone.

### RNA extraction and cDNA synthesis

mRNA isolation was performed with InnuPREP RNA mini kit (Analytikjena, cat. no. 845-KS-2040250, Jena, Germany) according to the supplier’s protocols. The quantity of RNA was measured on an Eppendorf BioPhotometer (Eppendorf, Hamburg, Germany) and considered pure if the absorbance at 260nm/280nm was more than 1.8. The concentration of 0.4μg RNA/10 μl of dH2O was used. cDNA synthesis was performed using High Capacity kit (Applied Biosystems, cat. No. 4368814, Foster City, CA, USA) according to the manufacturer’s instructions on a TGradient thermocycler (Biometra, Göttingen, Germany).

### PCR

Quantitative RT-PCR analysis was performed using 20 ng cDNA per reaction using the Taqman Universal PCR Master Mix (Thermo Fisher, cat. No. 4304437) and SYBR^®^ Green PCR Master Mix (Thermo Fisher, cat. No. 4344463). Gene expression values were calculated using the mean C_t_ values of the samples. The expression of target genes was normalized to the housekeeping gene ACT, and then to St-T1b cells line (2^−ΔΔCt^). The primers were synthesized by Thermo Fisher and are listed in tables 2 and 3.

**Table 2.**
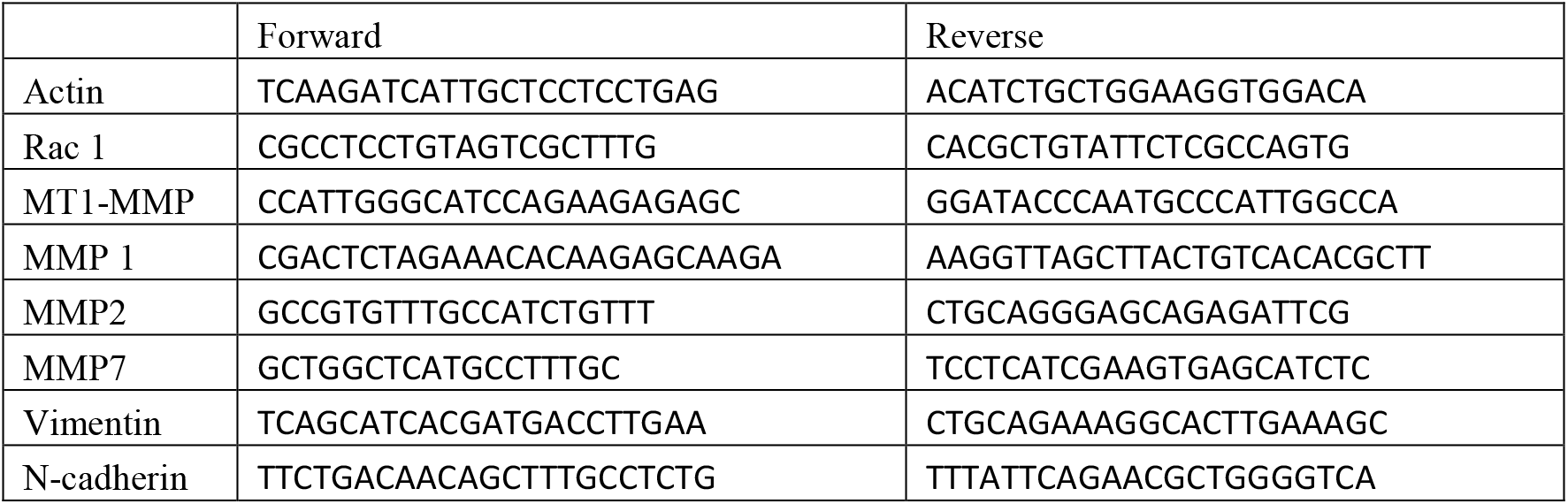
Sybr Green PCR Primers

**Table 3.** Taqman PCR Primers

|  |  |
| --- | --- |
| actin | hs99999903 m1 |
| ROCK2 | hs00153074 m1 |

### Statistical analysis

Data were analysed using GraphPad Prism8 (GraphPad Software, San Diego, USA). Normal distribution was tested using the Shapiro-Wilk test. A two-tailed unpaired Student’s t-tests was used to analyse statistical significance between two conditions in an experiment. For experiments with three or more comparisons, an ordinary one-way ANOVA with a Tukey’s multiple comparisons test was used. For data that were not normally distributed, the Kruskal-Wallis test followed by Dunn’s multiple comparisons test was used. A two-way repeated measures (RM) ANOVA with Šidák’s multiple comparisons test was used to evaluate the effect of Matrigel and collagen I on spheroid size over time. Significance values were chosen as *P<0.05; **P<0.01; ***P<0.001, ****P<0.0001. Error bars represent the mean±s.d or mean+s.d.. All figure panels were assembled in Inkscape 0.92.

## Author contributions statements

A.S. performed the majority of the experiments, drafted the figures and wrote the manuscript draft. V.F. performed experiments under the supervision of A.S. M.N., Y.S. and M.-K. v.W. helped to establish the 3D system by assisting with experiments, and by providing unpublished data and expertise in 3D culture. S.D.S and L.K. provided patient tissues and documented clinical data. B.G. provided resources, advice and expertise in 3D and primary cell culture and confocal immunofluorescence microscopy. L.K. provided resources and general support. M.G. and A.S conceived the study. All authors reviewed the manuscript.

## Acknowledgements

We would like to acknowledge Anna Starzinski-Powitz for the generous gift of 12Z cells, and Birgit Gellersen for the generous gift of the St-T1b cell line, Birgit Pers and Dorothea Godulla for expert technical assistance, Niki Loges for help with confocal microscopy, and Peter Friedl for helpful discussions. This research was supported by a WiRe – Women in Research Fellowship and a WWU Fellowship of the University of Münster (to AS) and European Commission (REA) EU H2020-MSCA-RISE-2015 grant 691058 MOMENDO (to MG). We acknowledge funding by the Open Access Publishing Fund of Münster University.

